# Spatial Scale of Non-Target Effects of Cotton Insecticides

**DOI:** 10.1101/2022.07.28.501895

**Authors:** Isadora Bordini, Steven E. Naranjo, Alfred Fournier, Peter C. Ellsworth

**Affiliations:** University of Arizona, Department of Entomology, Maricopa Agricultural Center, 37860 W Smith-Enke Rd, Maricopa, AZ 85138, USA; USDA-ARS, Arid-Land Agricultural Research Center, 21881 North Cardon Lane, Maricopa, AZ 85138, USA

## Abstract

Plot size is of practical importance in any integrated pest management (IPM) study that has a field component. Such studies need to be conducted at a scale relevant to species dynamics because their abundance and distribution in plots might vary according to plot size. An adequate plot size is especially important for researchers, technology providers and regulatory agencies in understanding effects of various insect control technologies on non-target arthropods. Plots that are too small might fail to detect potential harmful effects of these technologies due to arthropod movement and redistribution among plots, or from untreated areas and outside sources. The Arizona cotton system is heavily dependent on technologies for arthropod control, thus we conducted a 2-year replicated field experiment to estimate the optimal plot size for non-target arthropod studies in our system. Experimental treatments consisted of plot sizes and insecticides in a full factorial. We established three plot sizes that measured 144 m^2^, 324 m^2^ and 576 m^2^. For insecticides, we established an untreated check, a positive control insecticide with known negative effects on the arthropod community and a selective insecticide. We investigated how plot size impacts the estimation of treatment effects relative to community structure (27 taxa), community diversity, individual abundance, effect sizes, biological control function and success of arthropod taxa with a wide range of mobility, including *Collops* spp., *Orius tristicolor, Geocoris* spp., *Misumenops celer, Drapetis* nr. *divergens* and *Chrysoperla carnea*. The 144 m^2^ plots supported similar results for all parameters compared to larger plots, thus being sufficiently large to measure insecticidal effects on non-target arthropods in Arizona cotton. Though results might be system-specific, they point to a scale of testing that should be considered when developing any IPM guidelines, especially for systems that share a similar fauna of predators and pests.

## Introduction

Plot size is of practical importance for anyone designing experiments involving mobile species, regardless of the study purpose. Because natural arthropod communities range freely over large agricultural areas, observations from small areas could be quite different from the dynamics and patterns that occur in large areas. Thus, plots need to be as large as practically possible for experiments to be applicable to commercial scales (Jepson & Thacker, 1990; Prasifka et al., 2005; Macfadyen et al., 2014).

Choosing an adequate plot size is particularly important in non-target arthropod assessments involving insect control technologies e.g. insecticides and transgenic *Bt* crops. Several authors have demonstrated that the distribution and abundance of arthropods in plots varies according to plot size (Jepson & Thacker, 1990; Pullen et al., 1992; Duffield & Aebischer, 1994; Kennedy et al., 2001; Prasifka et al., 2005; Macfadyen et al., 2014). This variation can be especially problematic if plots are too small to detect harmful effects of these technologies on non-target arthropods due to their movement and redistribution among plots, or from untreated areas and outside sources (Macfadyen et al., 2014).

However, larger plots necessarily result in higher costs associated with land rental, water, labor and equipment, and might not be tenable when there is a limited amount of seed in the case of regulated trials. Plots need to be as large as practically possible while balancing the goals of accurately measuring gross effects on non-target arthropods and testing on an area suitable for the ecological attributes of the arthropod community, especially regarding mobility (Macfadyen et al., 2014; Alix et al., 2012).

Some authors have suggested that few studies have been conducted in field plots of sufficient size to understand potential disruptions of insecticidal technologies on biological control agents and other non-target organisms (Furlong & Zalucki, 2010, Zalucki et al., 2014, Macfadyen et al., 2014; 2015). Such studies need to be conducted at a scale that is relevant to species dynamics in any integrated pest management (IPM) study that has a field component, and to allow regulatory bodies to properly evaluate data submitted for insecticide or trait registration by technology providers. While harmful effects of transgenic crops on non-target arthropods are likely relatively small compared to insecticides (Naranjo, 2005), even minor issues with experimental design could compromise the ability to detect non-target effects (Wold et al., 2001; Prasifka et al., 2005).

We conducted a 2-year field experiment to estimate the optimal plot size for non-target arthropod studies in the Arizona cotton system. Our IPM program is heavily dependent on selective insecticides and the conserved biological control provided by key predators of our two key pests, *Lygus hesperus* Knight and whiteflies, *Bemisia argentifolii* Bellows and Perring (=*B. tabaci* MEAM1) (Ellsworth & Martinez-Carrillo, 2001; Naranjo and Ellsworth, 2009; Ellsworth et al., 2018; Vandervoet et al., 2018; Bordini et al., 2021). We investigated how plot size impacted our ability to measure treatment differences using multiple metrics including individual predator species abundance, arthropod community structure and diversity, and biological control function.

## Materials and Methods

### Experimental Design

Experiments were conducted at the University of Arizona’s Maricopa Agricultural Center, Maricopa, AZ, United States, in 2017 and 2018. Cotton, *Gossypium hirsutum* L., was planted on 1 June 2017 and 4 May 2018, and grown in accordance with agronomic practices for the area. The variety planted in both years, DP1549B2XF, was a Bollgard II® XtendFlex® variety (Bayer Company, MO, USA), which provides resistance to lepidopteran insects and tolerance to dicamba, glyphosate and glufosinate herbicides.

A randomized complete block design was used in both years. Plots were established in a single field site about 3 ha in size, subdivided into four blocks. Within blocks, treatments were randomly assigned to plots.

Experimental treatments consisted of plot sizes and insecticides in a full factorial. Three square plot sizes were established: a “small” plot: 12 m long by 12 m (12 rows; 144 m^2^); “medium” plot: 18 m long by 18 m (18 rows; 324 m^2^); “large” plot: 24 m long by 24 m (24 rows; 576 m^2^) with ca. 1 m row spacing, and 3 m unplanted alleys. For insecticides, two controls were established, a negative control, the untreated check (water only, UTC), and a positive control (a broad-spectrum insecticide) with known negative effects on the arthropod community (Bordini et al., 2021). This positive control enabled us to assess the ability of our experimental design to detect an expected effect on the arthropod community, and was implemented as acephate (Orthene® 97P, Amvac Chemical Corporation, California, USA) at 1120 g a.i./ha. Acephate has commercial activity against *L. hesperus*, but essentially no effect on *B. argentifolii*. A third treatment consisted of a selective insecticide that targets *B. argentifolii* and with proven non-effects on the arthropod community (Bordini et al., 2021). The selective insecticide was flupyradifurone (Sivanto™ 200SL, Bayer Crop Science, North Carolina, USA) applied at 202 g a.i./ha.

The trial was sprayed with a six row (6 m) tractor-mounted boom sprayer (TJ69-8003VS TeeJet® spray tips, two over the top nozzles per row) at a volume of 112.5 l/ha. To avoid drift, insecticides were sprayed directly to plots during calm weather conditions, using low spray boom heights and reduced sprayer ground speed.

Calendar sprays with acephate and flupyradifurone treatments were made every 14 days for a total of 3 sprays at their highest labeled rates during the flowering period. These were scheduled sprays for the purposes of non-target evaluations, and therefore they were not based on needs and/or thresholds for pest control. Spray dates were 8/1, 8/15 and 8/29 in 2017, and 8/2, 8/16 and 8/30 in 2018. A plant growth regulator (mepiquat pentaborate, Pentia™ 99 g ai / l, BASF, Texas, USA) was sprayed in 2017 according to cotton commercial guidelines to manage the balance between vegetative and reproductive growth for cotton production. These plant growth regulator sprays were not necessary in 2018.

### “Maintenance Sprays” for Prey Uniformity

Excess prey or prey resources can culminate in plot conditions more attractive (i.e., excess honeydew due to whiteflies), or less attractive (e.g., flower loss due to Lygus damage) to arthropods, causing gross changes across treatments that could potentially mask the effects of the intended treatments on non-target organisms (Bordini et al., 2021). The objective of these “maintenance sprays” was to preserve prey parity (*L. hesperus* and *B. argentifolii*) as much as possible among insecticidal treatments by spraying other insecticides that selectively targeted either whiteflies or Lygus (Bordini et al., 2021). Since insecticidal treatments provide control of different pests and one treatment is broad-spectrum (acephate controls *L. hesperus*) and the other one is selective (flupyradifurone controls *B. argentifolii*), we anticipated that there would be disparities among major arthropod prey, so we tried to maintain comparable levels of *B. argentifolii* and *L. hesperus* in all plots as much as possible with these “maintenance sprays”. The most convenient metric for those levels is the action threshold for each pest. Thus, these sprays targeted *L. hesperus* and *B. argentifolii*, and were deployed at economic threshold levels based on standard sampling methods for these pests (Ellsworth and Barkley, 2001, Ellsworth et al., 2006).

Maintenance sprays for *L. hesperus* control were deployed in the untreated check and flupyradifurone treatments, but not in the positive control as acephate has commercial activity against Lygus. The maintenance sprays for *L. hesperus* control were a selective insecticide (flonicamid, Carbine® 50WG, 98 g ai/ha, FMC Corporation, Pennsylvania, USA) applied twice in 2017 (10 and 24 August) and three times (2, 16, 30 August) in 2018. Maintenance sprays for *B. argentifolii* control were deployed once in the positive control and once in the untreated check in both years. We sprayed the selective insecticides, pyriproxyfen (Knack® 0.86EC, 75 g ai/ha, Valent, California, USA) on 24 August 2017, and buprofezin (Courier® 3.6SC, 390 g ai/ha, Nichino America, Delaware, USA) on 30 August 2018. As flupyradifurone has whitefly activity, no maintenance sprays were required against whiteflies; density levels there never exceeded the threshold.

### Arthropod Sampling

Lygus (nymphs and adults) and other arthropods were sampled concurrently with a standard 38 cm diameter sweep net. Sampling was done at three, seven and 13 days after spray (a total of 9 weekly dates over the season each year). Twenty-five sweeps per plot were used in the small plot, 2 sets of 25 sweeps (50 sweeps) in medium plots, and 4 sets of 25 sweeps (100 sweeps) in large plots. This intensity of sampling was similar across different plot sizes to ensure similar removal of arthropods. All data were standardized to 100 sweeps, because this is the unit of measurement used for *L. hesperus* and predator sampling in our system (Ellsworth and Barkley, 2001; Vandervoet et al. 2018).

Densities of *B. argentifolii* were sampled at three and seven days after each spray, for a total of six samples each year. Ten leaves from the fifth mainstem node below the terminal per plot were randomly selected to estimate adult density and collected to estimate egg and nymph density in the laboratory. Adult density was estimated by counting individuals on the underside of leaves *in situ* (Naranjo and Flint, 1995). Nymph and egg densities were estimated by counting individuals in the laboratory under magnification on a 3.88 cm^2^ disk taken from these leaves (Naranjo and Flint, 1994).

Densities of 27 additional arthropod taxa were estimated, including key arthropod predators in our system (Vandervoet et al., 2018); *Collops quadrimaculatus* (Fabricius), *C. vittatus* (Say), *Orius tristicolor* (White), *Geocoris punctipes* (Say), *G. pallens* Stål, *Misumenops celer* (Hentz), *Drapetis* nr. *divergens* Loew and *Chrysoperla carnea s.l*. (Stephens). Samples were frozen and later counted in the laboratory using a dissecting microscope.

### Biological Control Function

#### Predator to Prey Ratios

The key predators mentioned above were used to calculate predator to prey ratios as the quotient of each species per 100 sweeps to the number of *B. argentifolii* adults per leaf or large nymphs per leaf disc. We estimated eight predator to prey ratios as follow: *M. celer/B. argentifolii* adults, *M. celer*/*B. argentifolii* large nymphs, *D*. nr *divergens*/*B. argentifolii* adults, *D*. nr *divergens*/*B. argentifolii* large nymphs, *O. tristicolor*/*B. argentifolii* adults, *C. carnea* larvae/*B. argentifolii* adults, *Collops* spp./*B. argentifolii* large nymphs and *G. punctipes/B. argentifolii* large nymphs. For *Geocoris spp*., evaluations were done only with *G. punctipes* because *G. pallens* Stål densities were very low throughout the years of these trials. We chose these ratios because at certain levels they independently indicate functioning whitefly biological control in our system (Vandervoet et al., 2018, Ellsworth et al., 2019a, Ellsworth et al., 2019b; Bordini et al., 2021). We also calculated the proportion of time that these ratios were at or above functioning biological control over the season (Bordini et al., 2021).

#### Sentinel Prey

A novel *in situ* sentinel prey method was developed to measure biological control function along with other sources of in-field mortality for *B. argentifolii* eggs and fourth instar nymphs based on procedures developed for life tables of sessile insects (Naranjo & Ellsworth 2005; Naranjo & Ellsworth, 2017). Whiteflies are convenient and realistic as sentinel prey, because nymphs and eggs are immobile on leaves, abundant in the field, and natural enemies readily feed on them. They also are the most important pest subjected to biological control in our system. We used fourth instar nymphs because they 1) are easily seen in the field, 2) are the last instars prior to adult emergence and thus it is possible to distinguish successful emergence of adults from marks of mortality, 3) mortality is greatest during the fourth stadium, followed by mortality during the egg stage (Naranjo & Ellsworth, 2005), 4) insecticide thresholds in our system are based on large nymph numbers in addition to adults, and 5) the majority of our primary natural enemies feed on nymphs. We used eggs because this stage is subject to the second highest mortality level, and some of our key predators (*O. tristicolor*; *Geocoris spp*. and *C. carnea* larvae) feed on whitefly eggs (Naranjo & Ellsworth, 2005).

Three cohorts of at least 25 newly-laid live eggs (< 1 day old) per plot were established on 1, 16 and 29 August in 2018. The first two cohorts were established the same day of the sprays, and the third cohort was established the day before the last spray. Two cohorts of at least 25 newly molted fourth instar live nymphs, appearing flat and translucent, per plot were established on 2 August in three replicate plots, and 29 August in all plots in 2018. The first cohort was established the same day of the first spray, and the second cohort was established the day before the last spray.

Leaves with nymphs were collected 4–5 days and leaves with eggs at 7 days after establishment and taken to the laboratory for inspection. These intervals correspond with developmental times for these stages in a typical Arizona summer. Eggs and nymphs were examined to determine infield mortality sources using a dissecting scope in the laboratory. A single observer made all determinations within a single block. Mortality was recorded as due to predation, parasitism, inviability or eggs that failed to hatch, dislodgement and unknown. Dislodgement can be due to weather like rain and/or wind or to chewing predation by beetles mainly from *Collops* spp. or Coccinelidae. This type of predation usually removes all trace of the individual. However, in rare cases we could observe partial nymphal cadavers or egg pedicels still anchored on leaves (Naranjo and Ellsworth, 2005). Predation was mainly due to the sucking predators *Geocoris* spp., *Orius* spp. and *C. carnea* larvae as shown in previous studies (Naranjo and Ellsworth, 2005). These predators evacuate the content of nymphs and eggs, leaving a transparent nymph cuticle and egg chorion on the leaf.

Different factors that affect *B. argentifolii* mortality occur simultaneously and so there is no obvious temporal sequence of mortality. Thus, mortality from one factor can conceal the action of a previous factor (Naranjo and Ellsworth, 2005). To account for this and estimate mortality accurately, marginal mortality rates were calculated for each factor (Royama, 1981; Buonaccorsi & Elkinton, 1990; Elkinton et al., 1992; see Naranjo and Ellsworth, 2005 for formulas).

### Diversity Indices

Arthropods collected from sweep samples were used to calculate Species Richness (S), the Shannon-Wiener Diversity Index (H), the Effective Number of Species associated with the Shannon Diversity Index (ENS) and the Shannon Evenness Index (J). These metrics were calculated for the untreated check, the positive control acephate and flupyradifurone in each plot size.

### Statistical Analyses

We used a mixed-model, repeated measures analysis of variance (JMP® Pro 14.2, SAS Institute Inc., Cary, NC) to test for treatment differences (insecticide and plot size) affecting the abundance of the six key predators and key pests over the season in both years. We also used this model to test for treatment differences in diversity metrics. The model included fixed effects of plot size, insecticide, year and sampling date (repeated measure). Block and associated interaction terms were considered random effects. The covariance structure used was AR(1). We used the proportion of maximum scaling method (POMS), which transforms each scale (predators’ scale in each year) to a common metric running from 0 (=minimum possible) to 1 (=maximum possible) (Moeller, 2015), to minimize expected year effects in abundance of the predatory arthropods and to enable more consistent comparisons between years. To meet normality and homoscedasticity assumptions, the arthropod predator data were transformed by sqrt(x + 0.5). *B. argentifolii* and *L. hesperus* and abundance were transformed using ln(x + 1) and sqrt(x + 0.5), respectively. We compared the mean weekly abundance of the six key predators, pests, diversity metrics using Dunnett’s test within each year. The untreated check for large plot size was the standard.

Analyses were done for all sample dates after the first application of insecticides. Before insecticidal application, pre-counts of arthropod densities were not statistically different. To facilitate the visualization of patterns, we graphically represented pest and predator abundance using cumulative arthropod-days over the season using the trapezoidal rule (Ruppel, 1983).

We calculated the proportion of dates that each key predator to prey ratio was above functioning biological control levels in our system (Bordini et al., 2021). These data were analyzed using a mixed-model that included fixed effects of insecticide treatment and year; the block variable and associated interaction terms were entered as random effects (JMP® Pro 14.2, SAS Institute Inc., Cary, NC). The same model was used to test for treatment differences (insecticide and plot size) in marginal mortality for nymphs and eggs. In all cases, Dunnett’s test was used to compare results from treatments with the large plot untreated check.

We examined the main effects of plot size on the entire arthropod predator community through Principal Response Curves (PRC), a time-dependent, multivariate analysis that depicts arthropod community trends over time for each treatment relative to a control (Van den Brink and Ter Braak, 1998; 1999; Ter Braak and Smilauer, 1998; 2012). We examined the main plot size effects using the large plot size as the standard. PRCs use a distribution-free *F* type test based on sample permutation to test for statistical significance in patterns. Principal response analyses were done in CANOCO v5 (Microcomputer Power, Ithaca, NY, USA).

Finally, we estimated Hedge’s d effect sizes between means from arthropod abundance (key predators, *Lygus* spp. and *B. argentifolii*) in each insecticidal treatment (positive control and flupyradifurone) relative to means in the untreated check for each plot size.

## Results

### Individual predator abundance

Plot size and its interactions were not significant for key predator abundance (P > 0.05; Table 1; Fig 1.). Predator abundance in the flupyradifurone treatment was either statistically greater or not different from the UTC most of the time, being lower than the UTC in only two instances (once for *O. tristicolor* after the first spray in 2017, and once for *M. celer* after the third spray in 2018). The negative control, acephate, supported much lower predator densities as expected. The results from the negative control provide strong evidence that the experimental design was robust enough to measure the known destructive effect of this insecticide.

**Table 1.** Fixed effect F-values for mean arthropod predator abundance (per 100 sweeps) over two years. See Fig. 1 for plots of density over the season.

| Fixed Factors | DF | *M. celer* | *G. punctipes* | *O. tristicolor* | *Drapetis nr divergens* | *C. carnea* larvae | *Collops* spp. |
| --- | --- | --- | --- | --- | --- | --- | --- |
| Chemistry | 2, 54 | 28.32*** | 45.88*** | 3.09 | 3.39* | 27.1*** | 4.88* |
| Plot Size | 2, 54 | 0.76 | 1.22 | 2.09 | 0.54 | 1.88 | 0.02 |
| Year | 1, 54 | 3.64 | 5.92* | 4.12* | 39.58*** | 0.11 | 13.94*** |
| Week | 8, 384 | 4.88*** | 7.18*** | 19.54*** | 28.46*** | 5.59*** | 4.3*** |
| Chemistry*Plot Size | 4, 54 | 1.58 | 0.38 | 0.45 | 0.57 | 2.26 | 0.57 |
| Chemistry*Week | 16, 410 | 1.61 | 2.85*** | 1.45 | 0.76 | 3.21*** | 0.93 |
| Chemistry*Year | 2, 54 | 0.58 | 0.70 | 0.09 | 0.59 | 1.01 | 1.68 |
| Chemistry*Year*Week | 16, 410 | 1.93* | 1.13 | 2.27** | 1.18 | 2.47** | 0.54 |
| Plot Size*Year | 2, 54 | 0.91 | 0.09 | 0.65 | 0.07 | 0.58 | 0.03 |
| Plot Size*Week | 16, 410 | 0.66 | 0.72 | 1.48 | 1.03 | 0.98 | 0.95 |
| Plot Size*Year*Week | 16, 410 | 0.81 | 0.45 | 1.50 | 1.30 | 0.71 | 0.64 |
| Week*Year | 8, 384 | 2.81** | 2.67** | 12.8*** | 31.27*** | 6.52*** | 5.98*** |
| Chemistry*Plot Size*Week | 32, 422 | 1.02 | 0.84 | 1.30 | 0.58 | 0.96 | 1.14 |
| Chemistry*Plot Size*Year | 4, 54 | 0.24 | 0.05 | 0.66 | 0.43 | 1.09 | 0.71 |

Repeated-measures ANOVA, \* $P < 0.05$ ; \*\* $P < 0.01$ ; \*\*\* $P < 0.001$ . DF are approximated.

**Figure 1.**
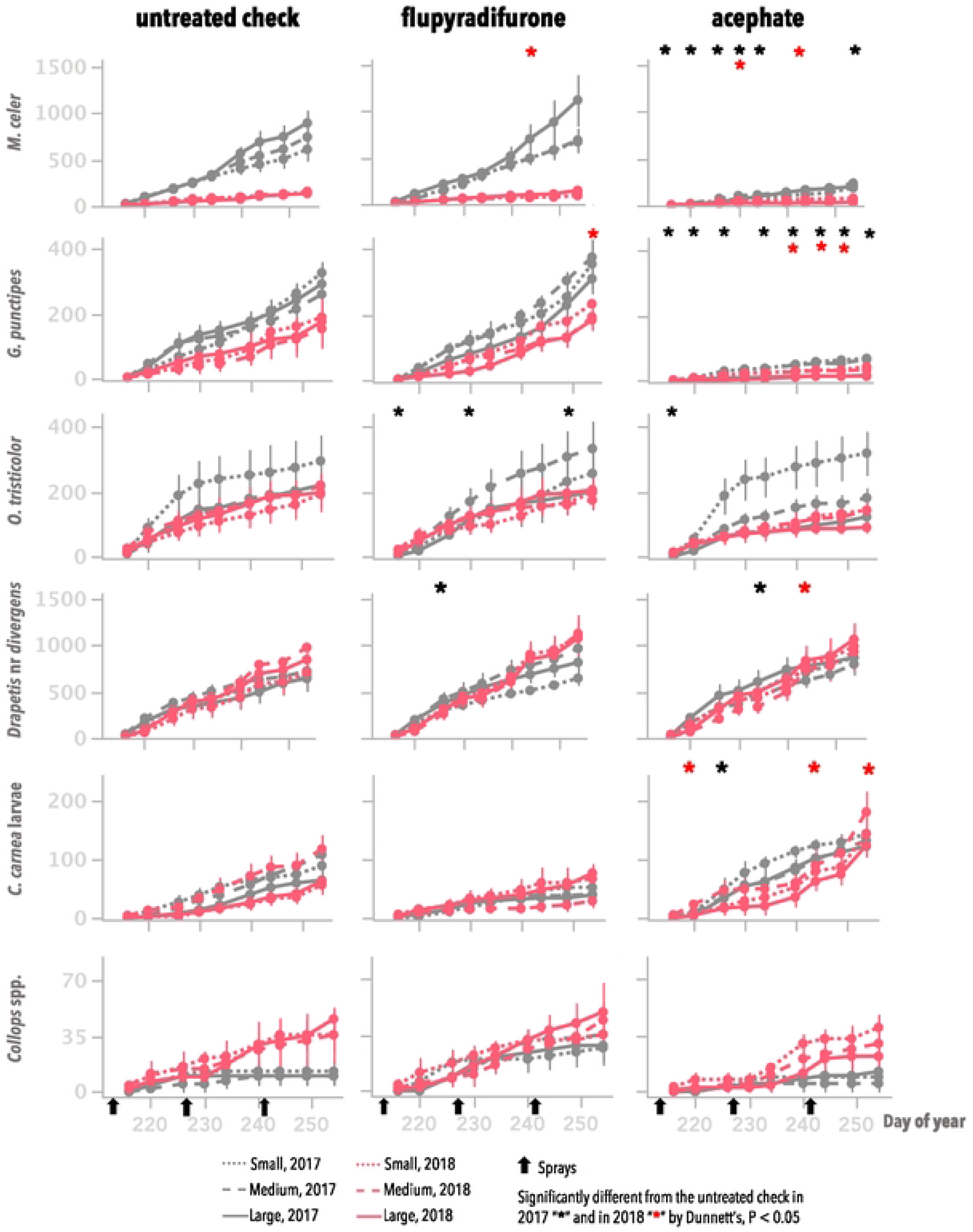
Post-treatment, cumulative mean insect-days (error bars = S.E) for arthropod predators per 100 sweeps during two growing seasons in Maricopa, AZ. Asterisks correspond to treatment means for the main effect of insecticides that were significantly different from the untreated check by Dunnett’s, P < 0.05, by week and year; plot size and its interactions were not significant.

### Non-target arthropod community dynamics

The Principal Response Curve (PRC) depicts the effect of small and medium plot sizes relative to the large plot size for the untreated check, positive control and flupyradifurone separately (Fig. 2). PRCs based on the first axis of redundancy analyses were not significant (P > 0.05) in any of the cases (Fig. 2). We also examined each insecticide compared to the untreated check for each plot size to further understand community effects (Fig S1-S4). As expected, the arthropod community was significantly reduced in the negative control relative to the UTC in both years and in all plot sizes (P < 0.05) (Fig. S1-S3). Again, this validates our experimental design because we were able to clearly measure the known destructive impacts of the negative control, acephate, on the arthropod community in all plot sizes.

**Figure 2.**
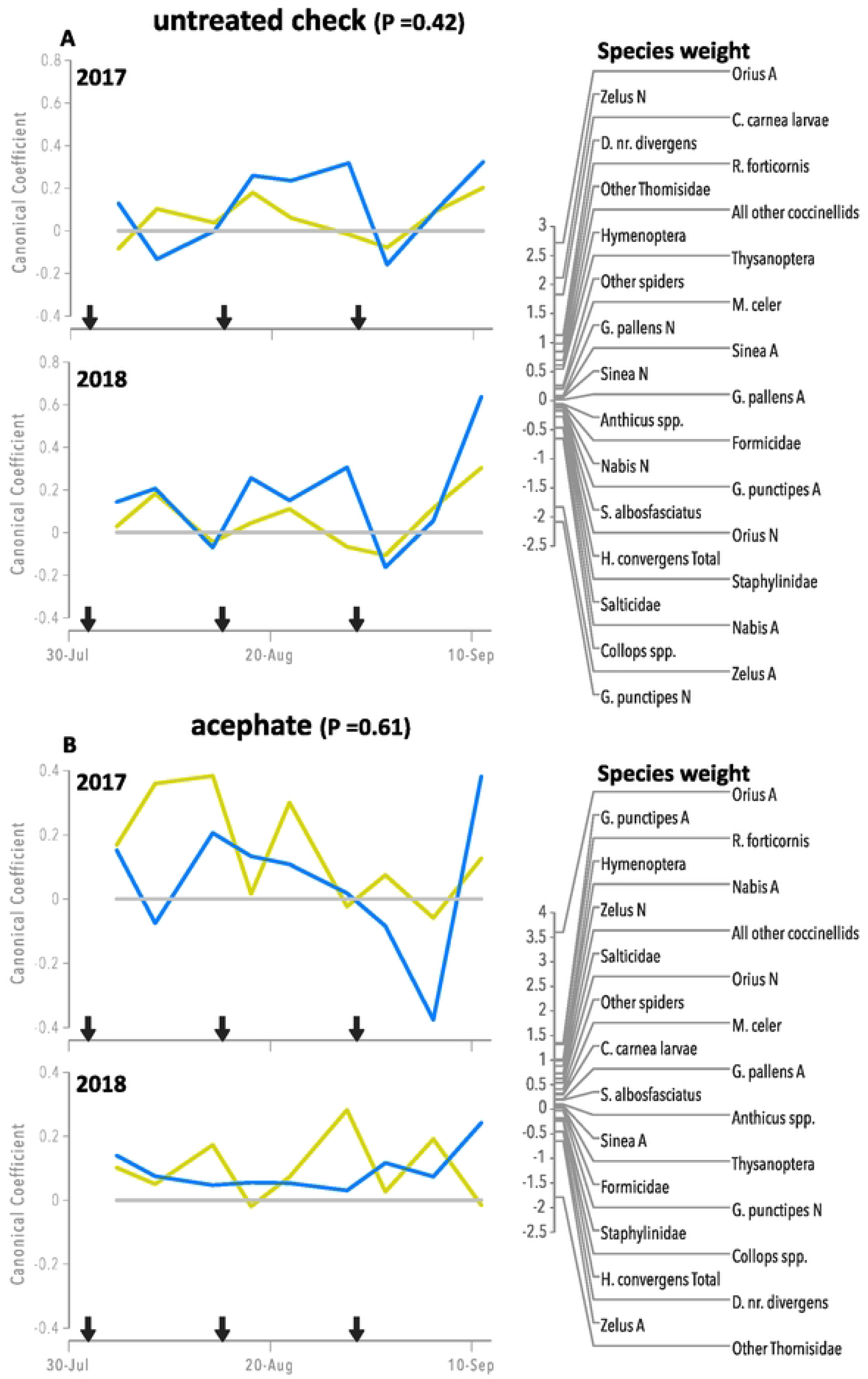
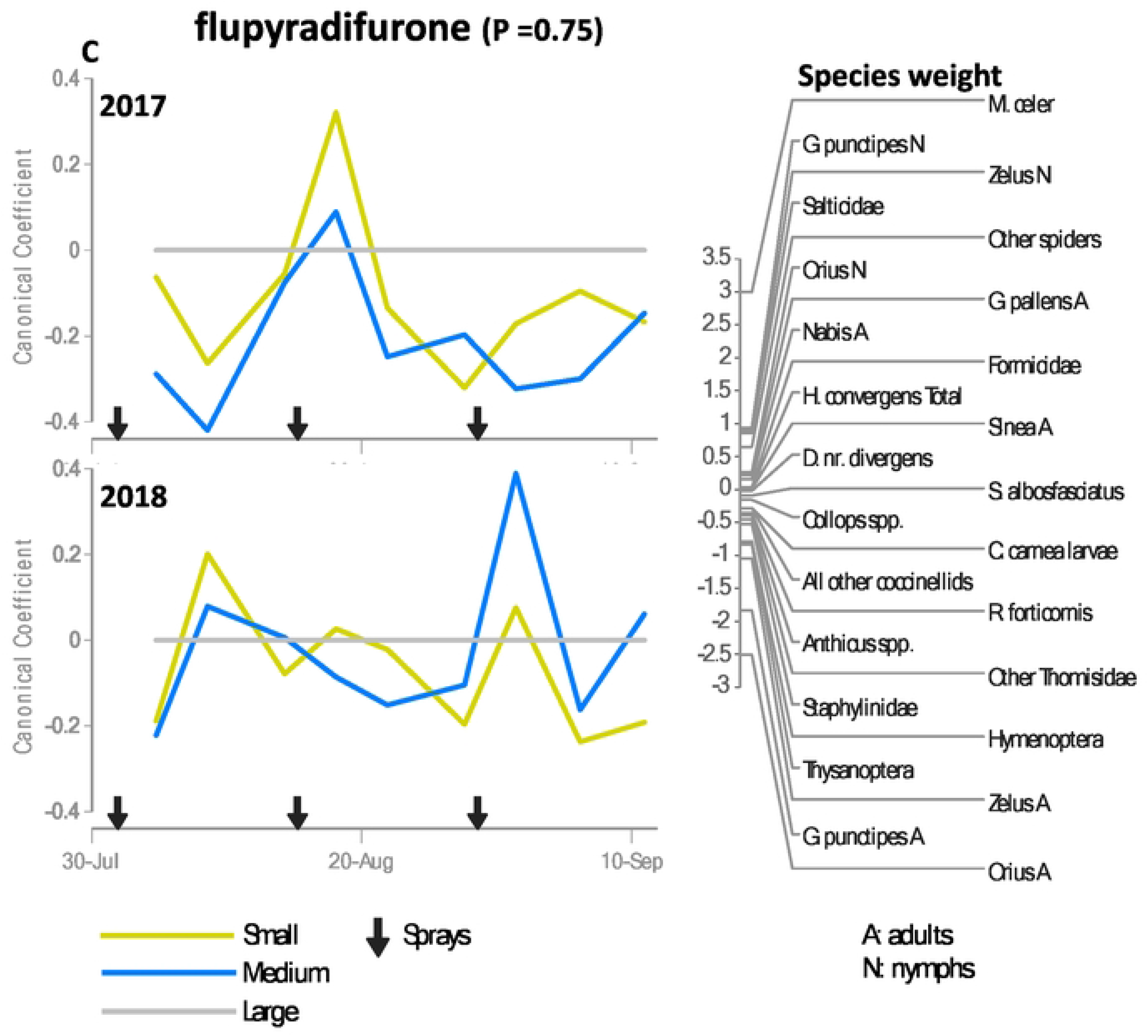
Principal response curves (PRC) showing plot size effects on the arthropod community for the untreated check (A), the positive control acephate (B) and flupyradifurone (C) relative to the large plot size (y = 0 line) during two growing seasons in Maricopa, AZ. The P-values denotes the significance of the PRC analysis in comparison with the large plot size over all dates based on an F-type permutation test. The product of the species weight and the canonical coefficient for a given plot size and time estimates the natural log change in density of that species relative to the large plot size. The greater the species weight the more the response for that species resembles the PRC. Negative weights indicate an opposite pattern, and weights between −0.5 and 0.5 indicate a weak response or a response unrelated to the PRC.

### Whitefly and Lygus target pest abundance

Plot size and its interactions were not significant for *B. argentifolii* and *L. hesperus* abundance (P > 0.05).

The abundance of *B. argentifolii* was significantly higher in the positive control than the UTC several instances in both years (Fig. 3; S.8). The abundance of *B. argentifolii* was significantly lower in the flupyradifurone treatment than the UTC in as many as four instances each year (Fig. 3; S.8).

**Figure 3.**
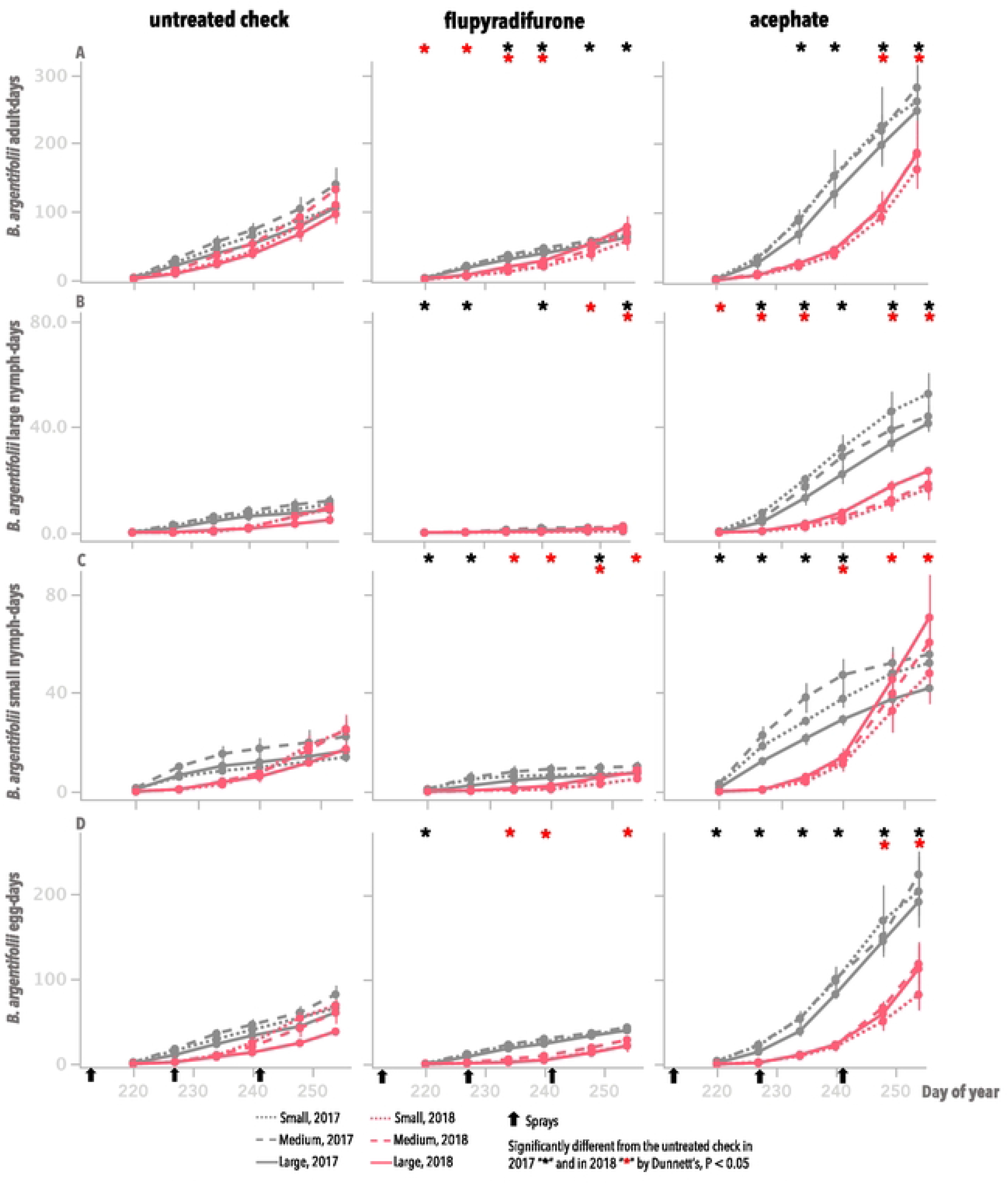
Cumulative mean insect-days (error bars = S.E.) for *B. argentifolii*, expressed as number of adults per leaf (A), large (B), small (C) nymphs and (D) eggs per 3.88 cm^2^ leaf disc. Asterisks correspond to treatment means for the main effect of insecticides that were significantly different from the untreated check by Dunnett’s, P < 0.05, by week and year; plot size and its interactions were not significant.

Population densities of *L. hesperus* were significantly lower in the positive control compared with the UTC on several dates in 2017 (mainly nymphs) and only in two dates in 2018 (Fig. 4). Population densities of this pest was significantly higher in the flupyradifurone than the UTC in only one date in 2017 (Fig. 4).

**Figure 4.**
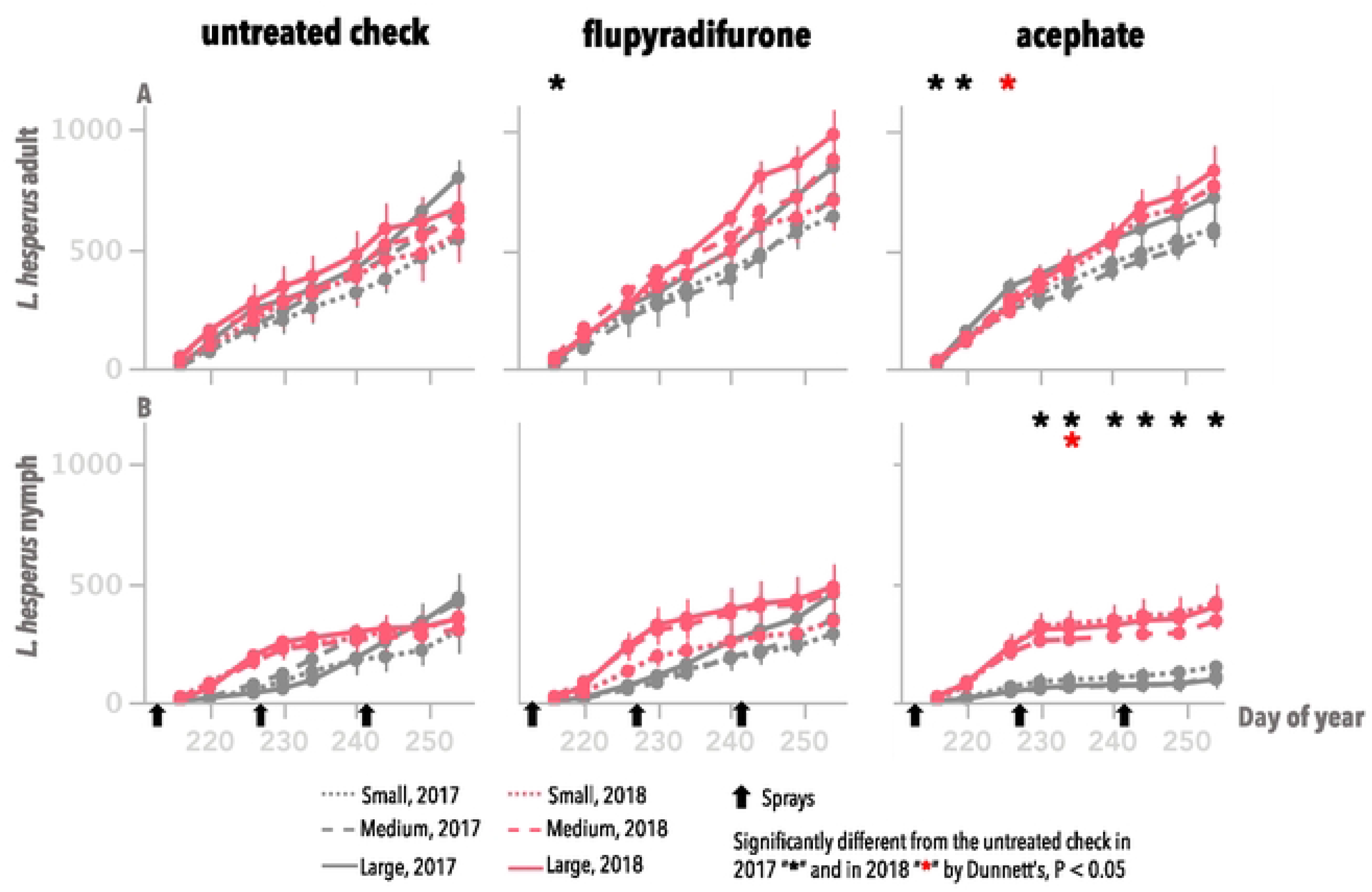
Cumulative mean insect-days for *Lygus hesperus* adults (A) and nymphs (B) per 100 sweeps. Asterisks correspond to treatment means for the main effect of insecticides that were significantly different from the untreated check by Dunnett’s, P < 0.05, by week and year; plot size and its interactions were not significant.

### Biological Control Function

#### Predator to Prey Ratios

We estimated the proportion of sample dates in which key predator to whitefly ratios were at or above levels indicative of functioning biological control in our system. Plot size and its interactions were not significant (P > 0.05; Table 2). The positive control performed as expected. The proportion of time at or above functioning biological control levels of multiple ratios were significantly lower in the positive control compared to the UTC (Fig. 5). This metric in the flupyradifurone treatment was significantly higher or not different from the UTC, with the exception of the ratio *C. carnea* larvae to *B. argentifolii* adult, in which the proportion of time was significantly lower than the UTC (Fig. 5).

**Table 2.** Fixed effect *F*-values of proportion of time that each of the eight predator to prey ratios were above levels associated with biological control of *B. argentifolii* in the Arizona cotton system.

| Fixed Factors | DF | *M. celer*/B. arg. nymph | *M. celer*/B. arg. adult | *D. nr divergens*/B. arg. nymph | *D. nr divergens*/B. arg. adult | *G. punctipes*/B. arg. nymph | *O. tricolor*/B. arg. adult | *C. carnea* larvae/B. arg. adult | *Collops spp.*/B. arg. nymph |
| --- | --- | --- | --- | --- | --- | --- | --- | --- | --- |
| Chemistry | 2, 54 | 31.76*** | 41.83*** | 69.17*** | 37.48*** | 57.15*** | 14.25*** | 11.84*** | 3.49* |
| Plot Size | 2, 54 | 0.04 | 0.27 | 0.68 | 0.98 | 1.19 | 0.25 | 0.08 | 0.43 |
| Chemistry*Plot Size | 4, 54 | 1.20 | 0.49 | 0.25 | 1.12 | 2.01 | 0.21 | 1.89 | 0.89 |
| Year | 1, 54 | 107.71*** | 101.01*** | 25.26*** | 24.49*** | 14.93*** | 0.33 | 0.11 | 7.37** |
| Chemistry*Year | 2, 54 | 2.20 | 3.61* | 8.26** | 2.13 | 0.21 | 1.08 | 1.33 | 0.76 |
| Plot Size*Year | 2, 54 | 0.28 | 0.09 | 0.30 | 3.00 | 0.86 | 0.36 | 1.11 | 0.27 |
| Chemistry*Plot Size*Year | 4, 54 | 0.87 | 0.19 | 0.77 | 0.61 | 0.47 | 0.82 | 0.77 | 1.46 |

Mixed-model ANOVA, \* $P < 0.05$ ; \*\* $P < 0.01$ ; \*\*\* $P < 0.001$

**Figure 5.**
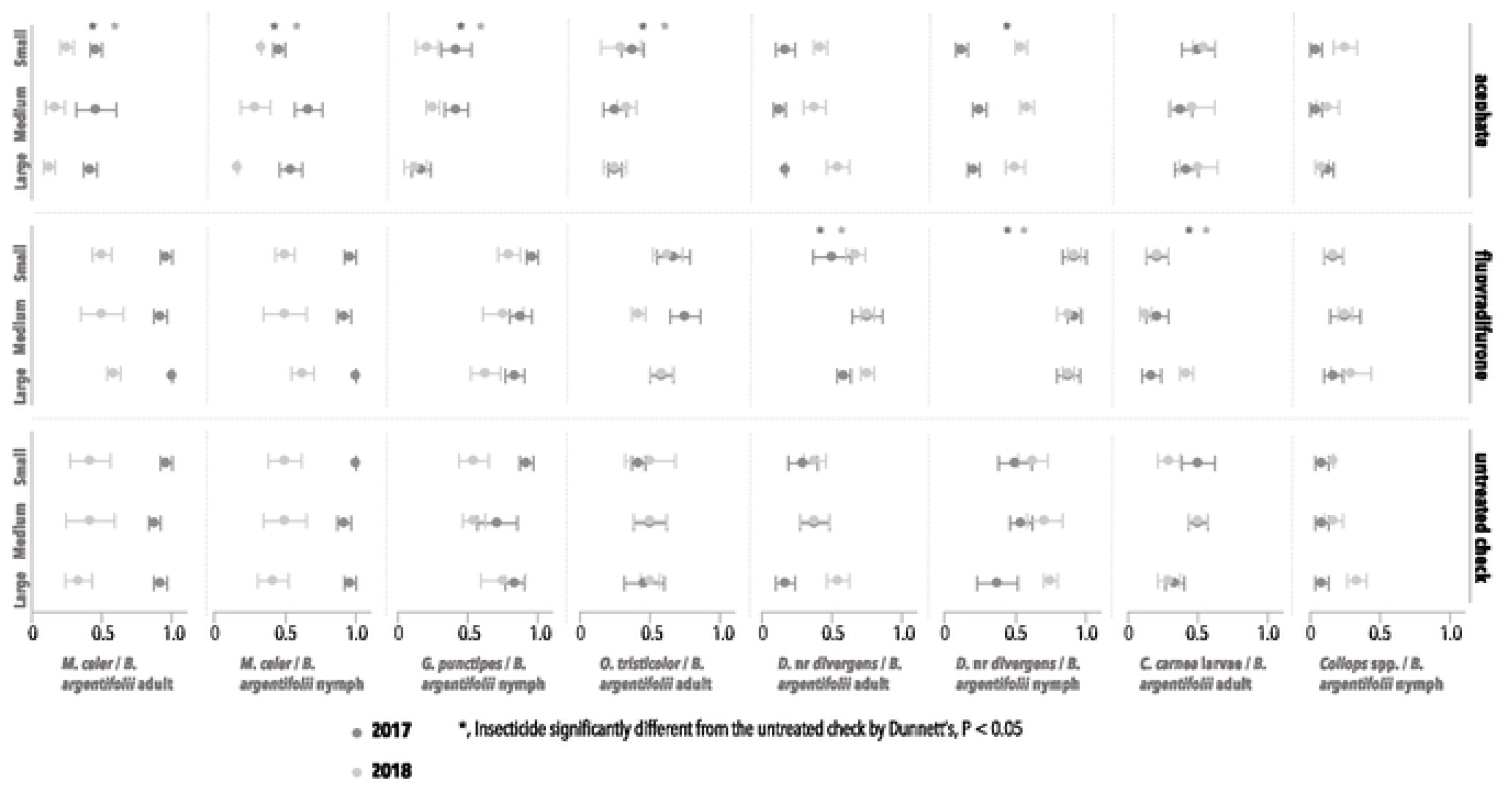
Proportion of time over the season that each of eight predator to prey ratios were above levels indicating functioning biological control (mean ± SE; Vandervoet et al., 2018) for each year. For each insecticide main effect, these proportions were compared with the UTC by Dunnett’s (*, P < 0.05) each year; plot size and its interactions were not significant.

#### Sentinel Prey

Neither plot size nor any of its interactions were significant for any mortality factor in the sentinel prey study with whitefly eggs or nymphs (P > 0.05; Tables 3 & 4). Total mortality, marginal predation and parasitism (nymphs only) of nymphs and eggs were significantly lower in the positive control than the UTC as expected (Fig. 6 & 7).

**Table 3.** Fixed effect *F*-values of mortality factors for *B. argentifolii* eggs.

| Fixed Factors | DF | Total mortality | Marginal predation | Marginal dislodgement | Marginal inviability |
| --- | --- | --- | --- | --- | --- |
| Date | 2, 49 | 9.74** | 3.6* | 41.52*** | 6.03** |
| Disruption | 1, 49 | 4.61* | 4.22* | 0.01 | 0.23 |
| Date*Disruption | 2, 49 | 2.15 | 3.01 | 0.22 | 1.78 |
| Plot size | 2, 49 | 1.10 | 1.62 | 0.17 | 0.16 |
| Date*Plot size | 4, 49 | 1.80 | 1.26 | 1.28 | 0.23 |
| Disruption*Plot size | 2, 49 | 0.26 | 0.09 | 0.18 | 0.91 |
| Date*Disruption*Plot size | 4, 49 | 1.96 | 1.82 | 1.14 | 0.14 |

Mixed-model ANOVA, \* $P < 0.05$ ; \*\* $P < 0.01$ ; \*\*\* $P < 0.001$

**Table 4.**
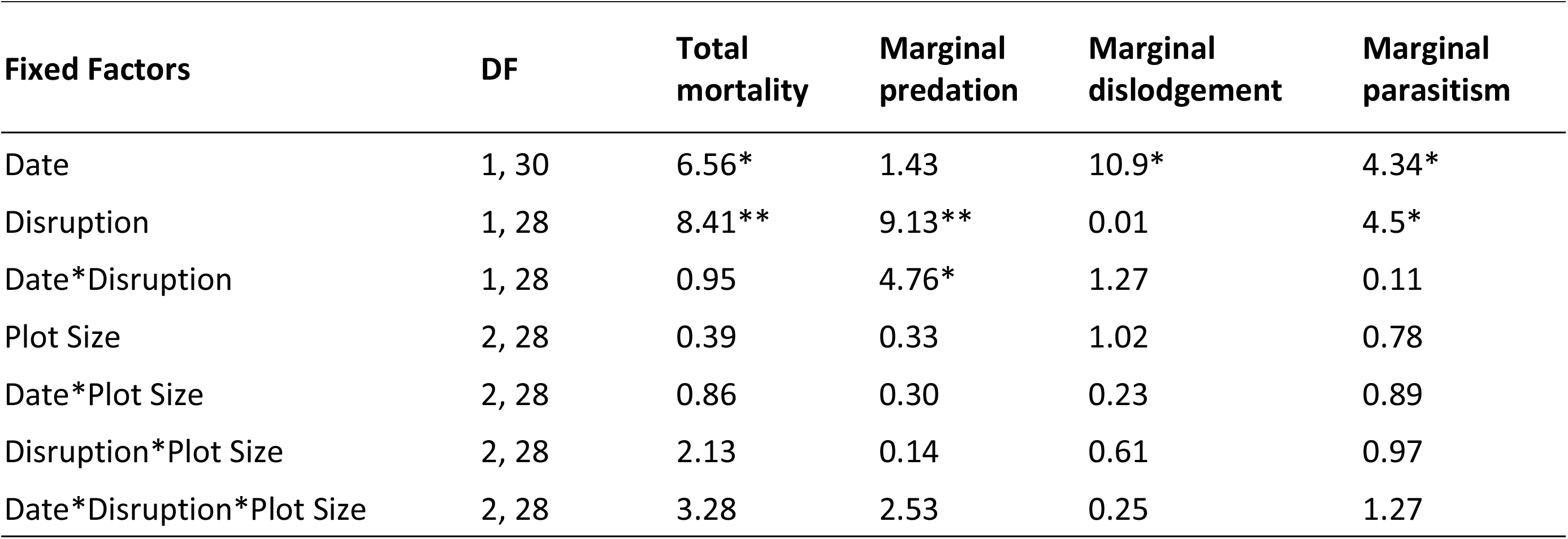
Fixed effect *F*-values of mortality factors for *B. argentifolii* nymphs.

| Fixed Factors | DF | Total mortality | Marginal predation | Marginal dislodgement | Marginal parasitism |
| --- | --- | --- | --- | --- | --- |
| Date | 1, 30 | 6.56* | 1.43 | 10.9* | 4.34* |
| Disruption | 1, 28 | 8.41** | 9.13** | 0.01 | 4.5* |
| Date*Disruption | 1, 28 | 0.95 | 4.76* | 1.27 | 0.11 |
| Plot Size | 2, 28 | 0.39 | 0.33 | 1.02 | 0.78 |
| Date*Plot Size | 2, 28 | 0.86 | 0.30 | 0.23 | 0.89 |
| Disruption*Plot Size | 2, 28 | 2.13 | 0.14 | 0.61 | 0.97 |
| Date*Disruption*Plot Size | 2, 28 | 3.28 | 2.53 | 0.25 | 1.27 |

**Figure 6.**
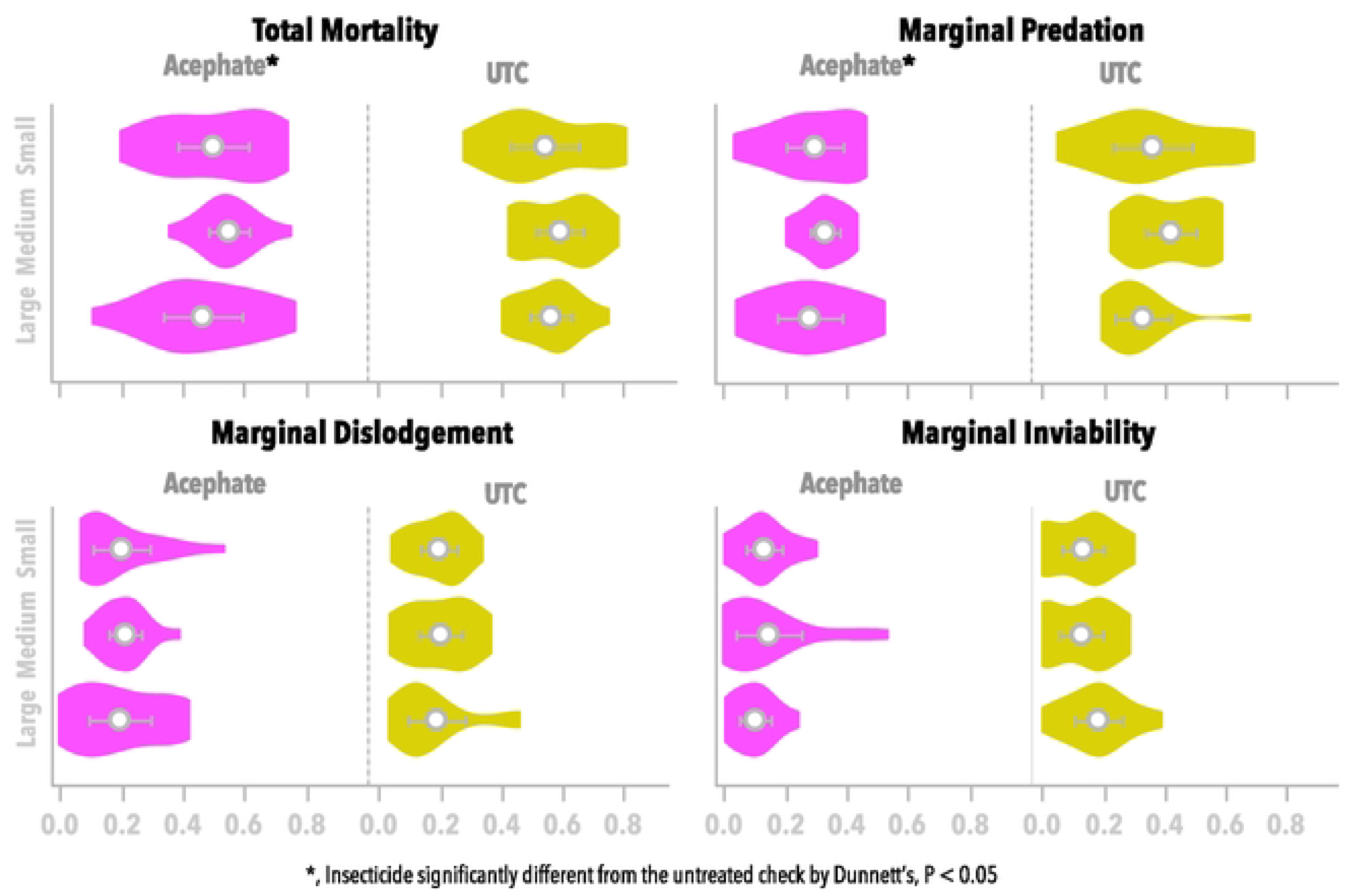
Violin plots showing distributions of mortality rates for *B. argentifolii* eggs (mean ± CI). Asterisks denote main effects for insecticides that were significantly different from the untreated check by Dunnett’s, P < 0.05; plot size and its interactions were not significant.

**Figure 7.**
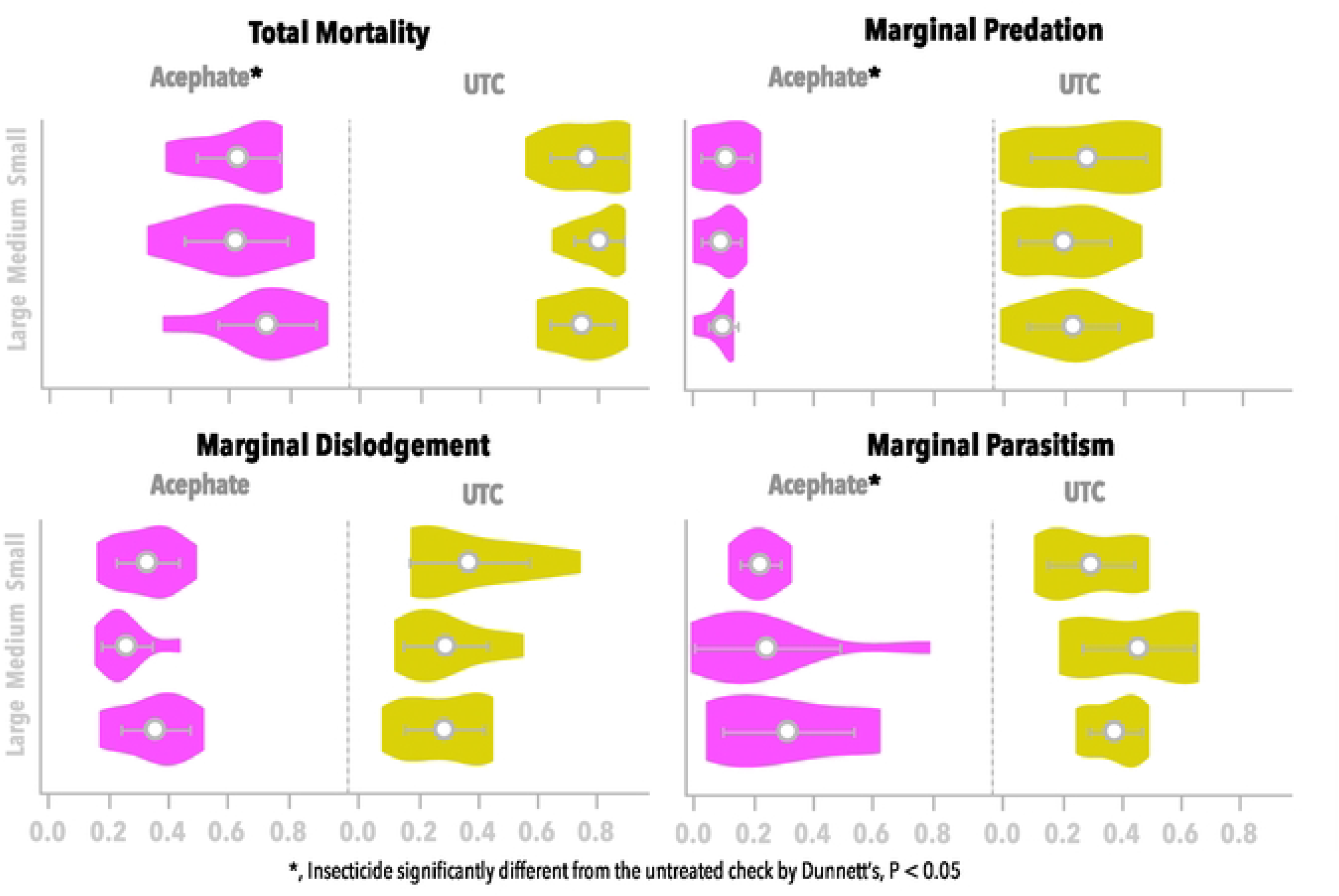
Violin plots showing distributions of rates for each mortality factor for *B. argentifolii* nymphs (mean ± CI). Asterisks correspond to treatment means for acephate that were significantly different from the untreated check by Dunnett’s, P < 0.05; plot size and its interactions were not significant. Marginal predation in the acephate treatment was significantly different from the untreated check only in the first nymph cohort by Dunnett’s, P < 0.05.

### Diversity Indices

None of the diversity indices were significantly affected by plot size or any of its interactions (P > 0.05; S5). There also were no strong patterns related to plot size for any of the indices (Fig. 8; S6).

**Figure 8.**
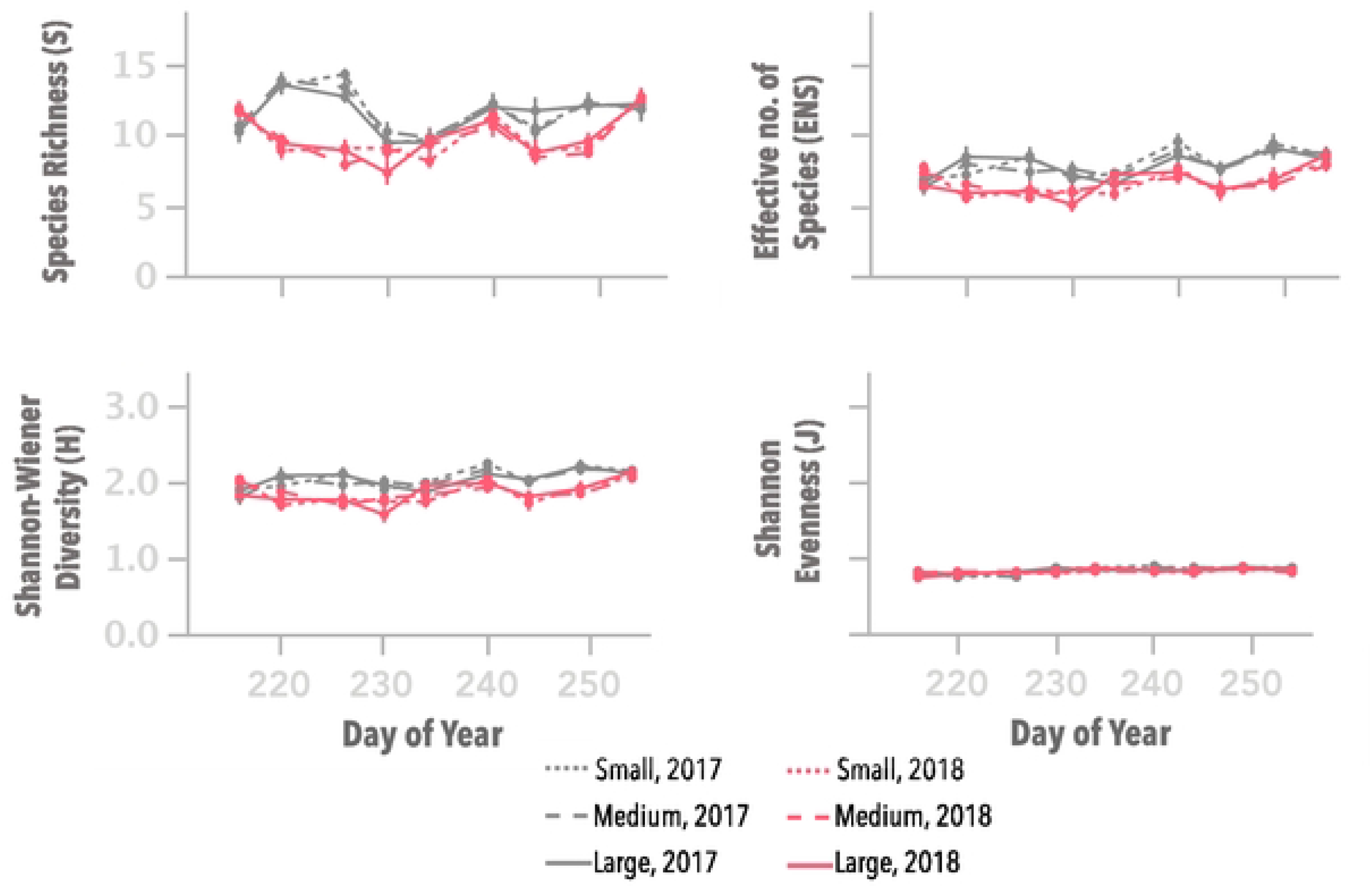
Post-treatment, main effect of plot size over all insecticide treatments (mean ± S.E) for diversity indexes during two growing seasons in Maricopa, AZ. Neither plot size nor any of its interactions were significant (P > 0.05).

### Effect Size

Effect size of predator abundance between the positive control acephate or flupyradifurone and the untreated check for each plot size ranged from small (0.2), medium (0.5) to large (0.8) effects sizes using Cohen’s scale (Cohen 1969). However, there were no patterns related with plot size (Table 5). We also estimated effect size for pest abundance, and there were again no patterns related with plot size (Table S7).

**Table 5.**
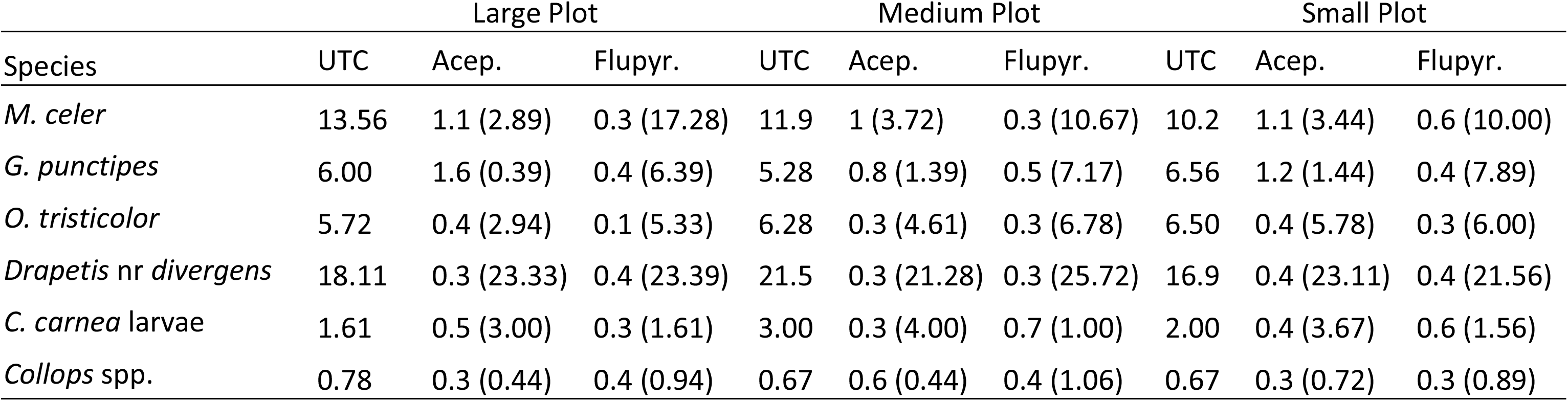
Effect size (density per 100 sweeps) between acephate or flupyradifurone and the untreated check for each plot size over two growing seasons. Only density is presented for the UTC.

| Species | Large Plot |  |  | Medium Plot |  |  | Small Plot |  |  |
| --- | --- | --- | --- | --- | --- | --- | --- | --- | --- |
|  | UTC | Acep. | Flupyr. | UTC | Acep. | Flupyr. | UTC | Acep. | Flupyr. |
| *M. celer* | 13.56 | 1.1 (2.89) | 0.3 (17.28) | 11.9 | 1 (3.72) | 0.3 (10.67) | 10.2 | 1.1 (3.44) | 0.6 (10.00) |
| *G. punctipes* | 6.00 | 1.6 (0.39) | 0.4 (6.39) | 5.28 | 0.8 (1.39) | 0.5 (7.17) | 6.56 | 1.2 (1.44) | 0.4 (7.89) |
| *O. tristicolor* | 5.72 | 0.4 (2.94) | 0.1 (5.33) | 6.28 | 0.3 (4.61) | 0.3 (6.78) | 6.50 | 0.4 (5.78) | 0.3 (6.00) |
| *Drapetis nr divergens* | 18.11 | 0.3 (23.33) | 0.4 (23.39) | 21.5 | 0.3 (21.28) | 0.3 (25.72) | 16.9 | 0.4 (23.11) | 0.4 (21.56) |
| *C. carnea* larvae | 1.61 | 0.5 (3.00) | 0.3 (1.61) | 3.00 | 0.3 (4.00) | 0.7 (1.00) | 2.00 | 0.4 (3.67) | 0.6 (1.56) |
| *Collops* spp. | 0.78 | 0.3 (0.44) | 0.4 (0.94) | 0.67 | 0.6 (0.44) | 0.4 (1.06) | 0.67 | 0.3 (0.72) | 0.3 (0.89) |

## Discussion

We investigated how plot size impacts the estimation of treatment effects relative to density, diversity and biological control function for arthropod taxa with a wide range of mobility (Arachnids, Coleoptera, Hemiptera, Diptera, etc.). We found no effect of plot size and concluded that “small” square plots (144 m^2^) are sufficiently large to measure insecticidal effects on nontarget arthropods in Arizona cotton. We were able to clearly measure the known destructive effects of the positive control (acephate), and detected the subtle effects including improved predation, and abundance of the selective insecticide flupyradifurone in small plots. Small plots supported arthropods for all parameters compared to medium and large plots based on several metrics: 1) individual species abundance and effect sizes of key predators and pests; 2) community structure (PRCs) and diversity (indices); and 3) biological control function (mortality of sentinel prey) and success (predator to prey ratios).

Previous plot size research done in Texas cotton found that “small” plots were sufficiently large for non-target arthropod studies. Harding et al. (1975) studied the effect of plot size on pests and predatory arthropods in cotton plots ranging from ca. 200 to 400 m^2^ treated with aerial sprays, and found that 200 m^2^ plots supported similar abundance of prey and predators compared to larger plots. Their 200 m^2^ plot was only three rows larger than our smallest plot size.

Despite our best efforts to control prey levels (whiteflies and Lygus) with “maintenance sprays” to achieve prey parity across insecticidal treatments, there were still significant differences in prey levels in some instances. Nonetheless, these significant differences in prey abundance were actually a strength in this study, because results from all parameters were consistent across plot sizes even at a variable abundance of prey among insecticidal treatments.

Plot size in non-target arthropod studies vary considerably (Macfadyen et al., 2014). For example, studies involving non-target species in arable crops used plots of 11,800 and 16,900 m^2^ in corn (Candolfi et al., 2004), and some recent studies used plots of 12 m^2^ in soybeans (Amin et al., 2022), 21 m^2^ in quinoa (Cruces et al., 2021) and 170 and 500 m^2^ in cotton (Marques et al., 2021; Kaur et al., 2021). In meta-analyses to examine the non-target impacts of transgenic Bt crops, Wolfenbarger et al. (2008) found no pattern in effect sizes between Bt and non-Bt crops over plot sizes ranging from 20 – 175,000 m^2^. This finding was recently supported by a comprehensive meta-analysis of Bt maize (Meissle et al. 2022). In contrast, studies conducted in cereals using a variety of plot sizes suggested that small plot trials may underestimate potential harmful effects of insecticides on non-target arthropods (Kennedy et al., 2001; Duffield & Aebischer, 1994; Pullen et al., 1992).

Our small plot here was the smallest practical plot that we could use without compromising sample size unit and over-sampling the fauna in our system. In our small plots, almost the entire length of two middle rows is needed to complete 25 sweeps and still avoid sampling the plot edges. It also was important to have a significant number of interior rows designated for sampling so that one could use alternate rows over time and minimize plant damage caused by sweeps. The minimal plot size is also a factor of the sampling method. Thus, systems that use other sampling methods (i.e., beat sheets) could potentially be subjected to smaller plots whereas other sampling methods might dictate the need for a larger plot. Plot geometry also needs to be considered. Our plots were square, however; long rectangular plots of the same area might not perform well because the distance between field edges of the longest side of rectangular plots would be reduced, which could influence arthropod movement and behavior differently.

At the other end of the spectrum, there were practical constraints to testing even larger plot sizes, including significant increases in costs of land rental, irrigation costs, labor, etc. Previous research done in larger plots in Arizona cotton found no effect of plot size in a long-term study testing the effects of transgenic Bt cotton on non-target arthropod abundance between plots ranging from 1200 m^2^ to 20,000 m^2^ (Naranjo, 2005). Larger plots are normally more heterogenous and might require greater sampling effort or more replicates to detect changes in population density (Naranjo, 2005). Another factor that needs to be considered is the potential loss of homogeneity of blocks as larger plots are used. Blocking field experiments control for nuisance variations like irrigation, soil texture, adjacent fields; however, at some point, inter-plot differences risk being larger than blocks differences or blocks cannot be efficiently established. Our results from the small plot size also suggest that interplot movement across bare ground was inconsequential because we were able to detect treatment differences in these small plots. Ground dwelling arthropods might behave differently, but they were not directly the subject of this investigation.

Our findings are both system-specific and general. They are system-specific in the sense of the arthropod fauna examined being specific to our system. On the other hand, we examined pests and predators that are found in other cotton systems across the country, and there are analog fauna in other regions globally. Thus, our results might be transferrable to locations that share a similar fauna and agriculture production system. Plot size might be influenced differently in other systems embedded in more diverse landscapes that impact surrounding areas and arthropod movement. Nevertheless, our approaches can be universally used in any system interested in investigating an optimal plot size for arthropod studies. Our results can be applied to any study in insect ecology in Arizona cotton, including studies involving transgenic cotton with insecticidal properties.

This study investigated the effect of plot size on several taxa with a wide range of mobility in Arizona cotton. We concluded that “small” square plots (144 m^2^) are sufficiently large for nontarget arthropod studies in Arizona cotton. Small plots allowed us to detect treatment differences, and supported similar individual predator abundance, arthropod community structure and diversity, and biological control function and success compared to larger plot sizes. Though results might be system-specific, they provide a baseline to choose a plot size sufficiently large to detect potential effects of insecticides and genetically modified crops in other systems, especially for those that share a similar fauna of predators and pests. This new information should be helpful to growers, researchers, technology providers and regulatory agencies in measuring impacts of various insect control technologies on non-target arthropods in cotton. Furthermore, they point to a scale of testing that should be considered when developing any IPM guidelines that are provided to farmers for use under commercial conditions. These results will guide Arizona’s field evaluation of current and future technologies with goals of providing reliable information about risk to non-target arthropods to our growers (Bordini et al., 2020).

## Acknowledgements

We thank F. Bojorquez, G. Lizarraga, J. Partida, A. Brown, E. Thacker, T. Thacker, N. Pier, P. Merten, M. Cruz, F. Pat, J. Fan and J. Trejo for their lab technical assistance. Thanks to P. Asiimwe for reviewing an earlier draft of this manuscript and providing helpful comments. We thank the pest control advisers Tom Montoya, Nathan Kempton, Ryan Tregaskes for supporting our proposal for this project. We thank the Entomology and Insect Science Graduate Interdisciplinary Program at University of Arizona. We also appreciate the funding support from the Western Integrated Pest Management Center, Western Sustainable Agriculture Research and Extension, Arizona Cotton Growers Association, United States Department of Agriculture – National Institute of Food and Agriculture (Extension Implementation Program, award number 2017-70006-27145) and Cotton Incorporated. Mention of trade names or commercial products in this article is solely for the purpose of providing specific information and does not imply recommendation or endorsement by the U.S. Department of Agriculture. USDA is an equal opportunity provider and employer.

## References

Alix A, Bakker F, Barrett K, Brühl CA, Coulson M, Hoy S, Jansen J-P, Jepson P, Lewis G, Neumann P, Sü ßenbach D and vanVliet. (2012). ESCORT 3: Linking non-target arthropod testing and risk assessment with protection goals. In Proceedings of the European Standard Characteristics Of non-target arthropod Regulatory Testing workshop ESCORT (Vol. 2).

Amin, Md, Sung-Dug Oh, Soo-Yun Park, Kihun Ha, Sera Kang, Jung-Ho Park, Minwook Kim, Chang Uk Eun, Young Kun Kim, and Sang Jae Suh. (2022). The effect of thioredoxin-gene-expressed transgenic soybean on associated non-target insects and arachnids. Plant Biotechnology Reports, 16(1), 79–90.

Bordini, I., Ellsworth, P. C., Naranjo, S. E., & Fournier, A. (2021). Novel insecticides and generalist predators support conservation biological control in cotton. Biological Control, 154: 104502.

Bordini, I., Fournier, A., Naranjo, S.E, Pier, N., Ellsworth, P.C. (2020). Cotton Insecticide Use Guide: Knowing and Balancing Risks. Arizona Pest Management Center field crops IPM shorts, University of Arizona, Cooperative of Extension. https://acis.cals.arizona.edu/docs/default-source/ipm-shorts/cottoninsecticiderisk.pdf?sfvrsn=a5628a8b_0 [accessed 20 April 2022].

Buonaccorsi, JP & Elkinton, JS. (1990) Estimation of contemporaneous mortality factors. Researches on Population Ecology 32: 151–171.

Candolfi, M. P., Brown, K., Grimm, C., Reber, B., & Schmidli, H. (2004). A faunistic approach to assess potential side-effects of genetically modified Bt-corn on non-target arthropods under field conditions. Biocontrol Science and Technology, 14(2), 129–170.

Cohen, J. (1969). Statistical Power Analysis for the Behavioral Sciences. Academic Press., New York, NY.

Cruces, L., de la Peña, E., & De Clercq, P. (2021). Field Evaluation of Cypermethrin, Imidacloprid, Teflubenzuron and Emamectin Benzoate against Pests of Quinoa (Chenopodium quinoa Willd.) and Their Side Effects on Non-Target Species. Plants, 10(9), 1788.

Duffield, S., Aebischer, N. (1994). The effect of spatial scale of treatment with dimethoate on invertebrate population recovery in winter wheat. Journal of Applied Ecology, 31(2), 263–281

Elkinton, J.S., Buonaccorsi, J.P., Bellows, T.S., Van Driesche, R.G., 1992. Marginal attack rate, k-values and density dependence in the analysis of contemporaneous mortality factors. Researches on Population Ecology 34, 29–44.

Ellsworth, P. C., & Martinez-Carrillo, J. L. (2001). IPM for Bemisia tabaci: a case study from North America. Crop protection, 20(9), 853–869

Ellsworth, P.C., Barkley, V. (2001). Cost-effective Lygus management in Arizona cotton. In: Silvertooth, J.C. (ed.), Cotton: A College of Agriculture and Life Sciences. University of Arizona, College of Agriculture and Life Sciences Tucson, AZ, pp. 299–307. https://repository.arizona.edu/handle/10150/211330 [accessed 4 April 2022].

Ellsworth, P.C., Fournier, A., Frisvold, G., Naranjo, S.E., 2018. Chronicling the socio- economic impact of integrating biological control, technology, and knowledge over 25 years of IPM in Arizona. In: Mason, P.G., Gillespie, D.R., Vincent, C. (Eds.), Proceedings of the 5th International Symposium on Biological Control of Arthropods. CAB International, pp. 214–216. https://acis.cals.arizona.edu/agricultural-ipm/vegetables/vegetable-outputs/publications/publications-view/chronicling-the-socio-economic-impact-of-integrating-biological-control-technology-and-knowledge-over-25-years-of-ipm-in-arizona

Ellsworth, P.C., Palumbo, J.C., Naranjo, S.E., Dennehy, T.J., Nichols, R.L. (2006). Whitefly management in Arizona Cotton 2006. The University of Arizona, Cooperative Extension, IPM Series No. 18. https://extension.arizona.edu/pubs/whitefly-management-arizona-cotton-2006 [accessed 4 June 2022].

Ellsworth, P.C., Pier, N., Fournier, A., Naranjo, S.E. (2019a). Making use of predators in cotton. Arizona Pest Management Center field crops IPM shorts, University of Arizona, Cooperative of Extension. https://cals.arizona.edu/crops/cotton/files/PtoPlaminate.pdf [accessed 4 April 2022].

Ellsworth, P.C., Pier, N., Fournier, A., Naranjo, S.E., Vandervoet, T. (2019b). Predator “thresholds”. Arizona Pest Management Center field crops IPM Shorts, University of Arizona, Cooperative of Extension. https://acis.cals.arizona.edu/docs/default-source/ipm-shorts/wfbit.pdf?sfvrsn=9e760083_2 [accessed 4 April 2022].

Furlong, M. and Zalucki, M. (2010). Exploiting predators for pest management: the need for sound ecological assessment. Entomologia Experimentalis et Applicata, 135(3), pp.225–236.

Harding, J. A., Dupnik, T.D., Wolfenbarger, D.A., & Fuchs, T.W. (1975). Evaluation of plot size for arthropod studies in cotton. Texas A&M Tech. Bull, 1–10.

Jepson, P., Thacker, J. (1990). Analysis of the spatial component of pesticide side-effects on non-target invertebrate populations and its relevance to hazard analysis. Functional Ecology, 4(3), p.349.

Kaur, J., Aggarwal, N., & Kular, J. S. (2021). Abundance and diversity of arthropods in transgenic Bt and non-Bt cotton fields under Indian conditions. Phytoparasitica, 49(1), 61–72.

Kennedy, P., Conrad, K., Perry, J., Powell, D., Aegerter, J., Todd, A., Walters, K. and Powell, W. (2001). Comparison of two field-scale approaches for the study of effects of insecticides on polyphagous predators in cereals. Applied Soil Ecology, 17(3), pp.253–266.

Macfadyen, S., Banks, J., Stark, J. and Davies, A. (2014). Using semifield studies to examine the effects of pesticides on mobile terrestrial invertebrates. Annual Review of Entomology, 59(1), pp.383–404.

Macfadyen, S., Davies, A., Zalucki, M. (2015). Assessing the impact of arthropod natural enemies on crop pests at the field scale. Insect Science, 22(1), pp.20–34.

Marques, L. H., B. A. Castro, J. Rossetto, O. A. Silva, V. F. Moscardini, L. H. Zobiole, A. C. Santos, P. Valverde-Garcia, J. M. Babcock, D. M. Rule, et al. (2021). Field efficacy of Bt cotton containing events DAS-21023-5× DAS-24236-5× SYN-IR102-7 against lepidopteran pests and impact on the non-target arthropod community in Brazil. PloS one, 16(5), e0251134.

Meissle, M., Naranjo, S. E., & Romeis, J. (2022). Does the growing of Bt maize change abundance or ecological function of non-target animals compared to the growing of non-GM maize? A systematic review. Environmental Evidence, 11(1), 1–36.

Moeller, J. (2015). A word on standardization in longitudinal studies: don’t. Front. Psychol. 6, 1389.

Naranjo, & Ellsworth, P. C. (2005). Mortality dynamics and population regulation in Bemisia tabaci. Entomologia Experimentalis et Applicata, 116(2), 93–108.

Naranjo, S. E. (2005). Long-term assessment of the effects of transgenic Bt cotton on the abundance of nontarget arthropod natural enemies. Environmental entomology, 34(5), 1193–1210.

Naranjo, S. E., & Ellsworth, P. C. (2009). Fifty years of the integrated control concept: moving the model and implementation forward in Arizona. Pest Management Science, 65(12), 1267–1286.

Naranjo, S., Ellsworth, P. (2017). Methodology for developing life tables for sessile insects in the field using the whitefly, Bemisia tabaci, in cotton as a model system. J. Visualized Experiments (129), e56150, doi:10.3791/56150.

Naranjo, S.E., Flint, H.M. (1994). Spatial distribution of preimaginal Bemisia tabaci (Homoptera, Aleyrodidae) in cotton and development of fixed-precision sequential sampling plans. Environ. Entomol. 23, 254–266.

Naranjo, S.E., Flint, H.M. (1995). Spatial distribution of adult Bemisia tabaci (Homoptera: Aleyrodidae) in cotton and development and validation of fixed-precision sampling plans for estimating population density. Environ. Entomol. 24, 261–270.

Prasifka, J., Hellmich, R., Dively, G., Lewis, L. (2005). Assessing the effects of pest management on nontarget arthropods: The influence of plot size and isolation. Environmental Entomology, 34(5), pp.1181–1192.

Pullen, A. J., Jepson, P. C., & Sotherton, N. W. (1992). Terrestrial non-target invertebrates and the autumn application of synthetic pyrethroids: experimental methodology and the trade-off between replication and plot size. Archives of Environmental Contamination and Toxicology, 23(2), 246–258

Royama, T., 1981. Evaluation of mortality factors in insect life table analysis. Ecological Monographs 51, 495–505.

Ruppel, R.F. (1983). Cumulative insect-days as an index of crop protection. J. Econ. Entomol. 76 (2), 375–377.

Ter Braak, C.J.F., Smilauer, P. (2012). CANOCO Reference manual and user’s guide: software for ordination (Version 5.0). Microcomputer Power, Ithaca, NY.

Ter Braak, C.J.F., Smilauer, P., (1998). CANOCO Reference Manual and User’s Guide to Canoco for Windows: Software for Canonical Community Ordination (Version 4). Microcomputer Power, Ithaca, NY.

Van den Brink, P.J., Ter Braak, C.J.F. (1998). Multivariate analysis of stress in experimental ecosystems by Principal Response Curves and similarity analysis. Aquatic Ecol. 32, 163–178.

Van den Brink, P.J., Ter Braak, C.J.F. (1999). Principal response curves: analysis of time-dependent multivariate responses of biological communities to stress. Environ. Toxicol. Chem. 18, 138–148.

Vandervoet, T.F., Ellsworth, P.C., Carriere, Y., Naranjo, S.E. (2018). Quantifying conservation biological control for management of Bemisia tabaci (Hemiptera: Aleyrodidae) in cotton. J. Econ. Entomol. 111, 1056–1068.

Wold, S. J., Burkness, E. C., Hutchison, W. D., & Venette, R. C. (2001). In-field monitoring of beneficial insect populations in transgenic corn expressing a Bacillus thuringiensis toxin. Journal of Entomological Science, 36(2), 177–187.

Wolfenbarger, L.L., Naranjo, S.E., Lundgren, J.G., Bitzer, R.J., Watrud, L.S., 2008. Bt crops effects on functional guilds of non-target arthropods: A meta-analysis. PLoS ONE 3, e2118.

Zalucki, M., Furlong, M., Schellhorn, N., Macfadyen, S., Davies, A. (2014). Assessing the impact of natural enemies in agroecosystems: toward “real” IPM or in quest of the Holy Grail?. Insect Science, 22(1), pp.1–5.

